# AI-based structural modelling and pathogenic variant mapping across the human protein interactome

**DOI:** 10.64898/2026.10.28.762585

**Authors:** Alexander M. Ille, Anetta L. Ille, Hannah H. Lamont, Todd Holyoak

## Abstract

Spanning the complex landscape of the human proteome, physical interactions between proteins coordinate cellular organization, signaling, regulation, and homeostasis. Yet the three-dimensional structures of most of these interactions remain unresolved. Here, we performed AI-based modelling of 109,282 human protein interaction structures, generating a structural resource for large-scale analysis. The predicted structures were systematically characterized through analysis of interface residue involvement, buried surface area, intermolecular bonding, and energetic properties. Additionally, we mapped pathogenic variant data onto the predicted structures, identifying interface-localized mutations for over 4,000 annotated disease categories. Acute myeloid leukemia, Alzheimer’s disease, and cardiomyopathy were examined as representative disease contexts for exploring the distribution of interface mutations within disease-associated protein interaction networks. Strengths and limitations of predictive capabilities were also assessed, including through experimental structure comparisons. Collectively, this structural resource and accompanying analyses offer a basis for developing novel hypotheses and informing experimental studies involving human protein interactions in physiological and disease contexts.

## Introduction

The organization and function of human cells depend on a diverse range of physical interactions across the proteome, which is encoded by ∼19,500 protein-coding genes [1]. Protein interactions mediate a broad range of biological processes, including gene expression, signal transduction, genome maintenance, and protein homeostasis, among others, with certain perturbations these interactions contributing to disease [2, 3]. Collectively, these interactions form the protein interactome, for which size estimates in humans span a broad, approximate range of 130,000-650,000 interactions [2, 4, 5]. Various databases have been continuously expanded to provide curated repositories of experimentally identified protein interactions [6-8]. In particular, the Biological General Repository for Interaction Datasets (BioGRID) [6, 9] was used as a source of protein interaction data for the current work.

Although human protein interactions have been extensively documented at the level of interacting partners, the three-dimensional structures of the vast majority of these interactions remain experimentally unresolved [10]. Experimental methods, primarily X-ray crystallography, cryogenic electron microscopy (cryo-EM), and nuclear magnetic resonance (NMR) spectroscopy, remain the gold-standard for the structural characterization of proteins and their interactions. However, the technical complexity, time, and resources required by experimental approaches make structural determination at the interactome scale currently impractical. Recent advances in artificial intelligence/machine learning (AI/ML) approaches for protein structure prediction [11-14] have been applied to address this gap in structural coverage of the human interactome [15-18]. The increasing availability of protein interaction structures predicted using various AI/ML-based approaches, despite inherent limitations and uncertainties, provides opportunities to advance our understanding of protein function and disease mechanisms on a large scale.

In the present study, we performed AI/ML-based structural modelling of 109,282 protein interactions and conducted a systematic characterization of the predicted structures. Protein interaction data and corresponding sequence data from BioGRID [6] and UniProt [19], respectively, were used as input for structural prediction using Boltz-2 [14], a diffusion-based AI/ML model with favorable performance in predicting protein interaction structures [17, 20]. We characterized the predicted structures through analyses of surface burial, residue interface involvement, intermolecular bonding, and energetic properties. Additionally, we mapped clinically-relevant pathogenic mutations sourced from the NCBI ClinVar database [21, 22] onto the interfaces of the predicted structures, identifying interface-localized mutational amino acid residues for over 4,000 annotated disease categories. Focusing on acute myeloid leukemia, Alzheimer’s disease, and cardiomyopathy as example disease contexts, we explored structurally-informed connectivity among protein interactions involving interface-localized mutations. We also performed comparisons with experimentally determined interaction structures obtained from the Protein Data Bank as well as identified various limitations and opportunities for interaction structure prediction improvement. Collectively, the predicted structures and accompanying analyses are intended to serve as a resource for generating hypotheses and informing studies on human protein interactions in physiological and disease contexts.

## Results

### Large-scale structural modelling of human protein interactions

We performed structural modelling for 109,282 experimentally annotated binary human protein interactions, generating a large-scale collection of *de novo* predicted three-dimensional interaction structures. The underlying protein interaction data were sourced from the Biological General Repository for Interaction Datasets (BioGRID) [6], a curated interaction resource in which individual records specify an interactor protein A and interactor protein B (a designation preserved in the current work). BioGRID data were pre-filtered to exclusively retain human-only, non-redundant interactions with direct contact between interacting protein pairs, and corresponding amino acid sequences were subsequently obtained from the UniProt Knowledgebase [19] (see Methods for details) **(Supplementary Table 1)**. The paired protein sequences were used as input for Boltz-2 [14] to jointly predict the three-dimensional structures of each protein interaction. An overview of the interaction structure prediction workflow is provided in **Fig. 1a**.

**Fig. 1:**
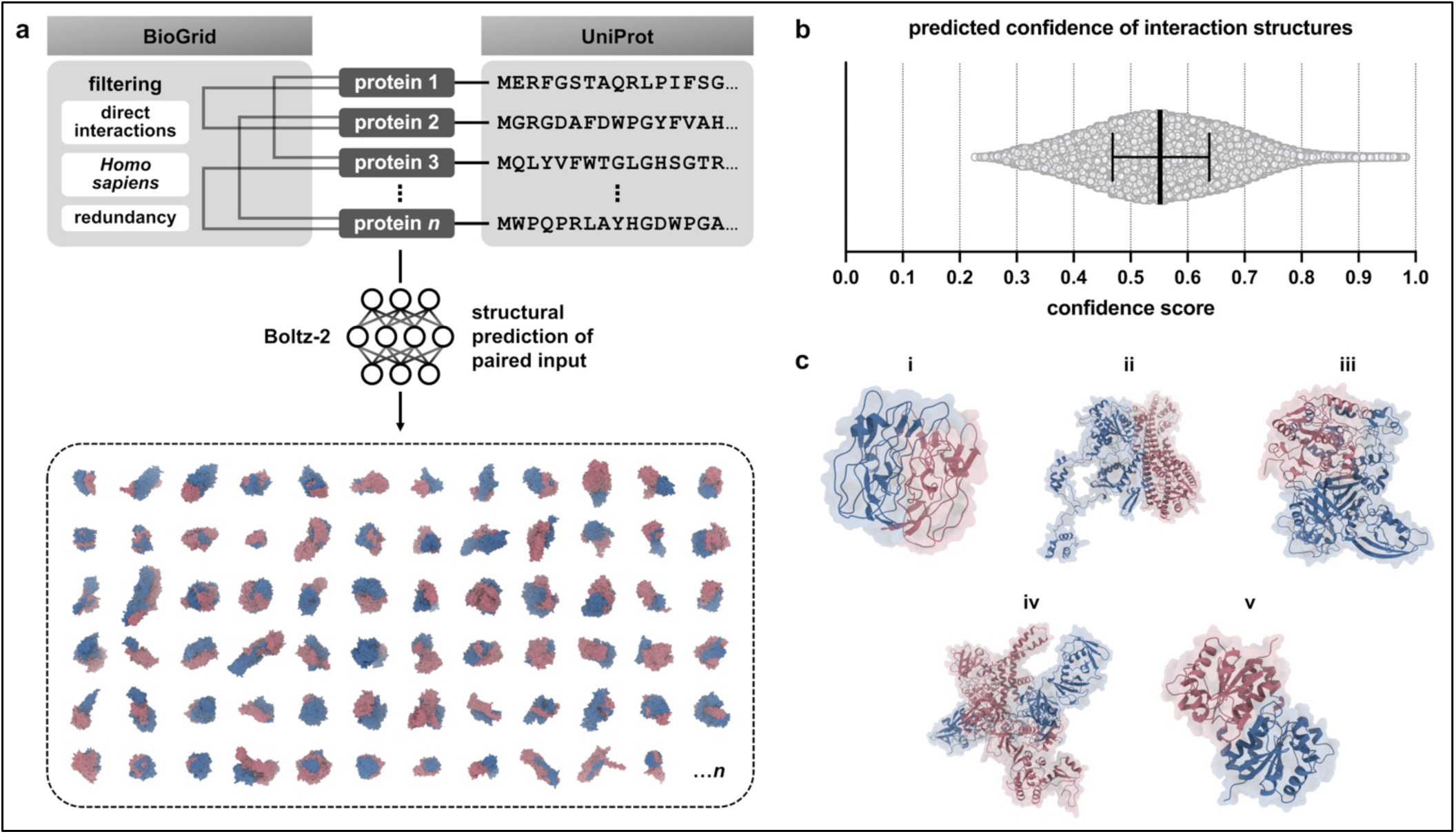
Overview of large-scale protein interaction structure prediction. **a.** Binary protein-protein interaction data from BioGrid was filtered to retain direct-only, human-only, and non-redundant interaction pairs. Corresponding sequences of these interaction pairs were obtained from UniProt. These paired sequences served as input for structural prediction using Boltz-2 for computing a total of *n* = 109,282 binary protein interaction structures, illustrated here with interactor protein A in red interactor protein B in blue. **b.** Distribution of confidence scores across predicted protein interaction structures, with each point representing one interaction structure. The median confidence score (0.5516) and interquartile range are shown; *n* = 109,282 interaction structures. **c.** Representative interaction structures of various confidence scores, including: (i) keratin-associated protein 11-1 (UniProt Q8IUC1) homomer, corresponding to the lowest confidence score (0.2272); (ii) DNA repair protein Ku70 (UniProt P12956) with uncharacterized protein KIAA0408 (UniProt Q6ZU52), corresponding to the lower Q1-adjacent confidence score (0.4685); (iii) E3 ubiquitin-protein ligase LNX (UniProt Q8TBB1) with josephin-1 (UniProt Q15040), corresponding to the median confidence score (0.5516); polyubiquitin-C (UniProt P0CG48) with peripheral plasma membrane protein CASK (UniProt O14936), corresponding to the lower Q3-adjacent confidence score (0.6381); and Parkinson disease protein 7 (UniProt Q99497) homomer, corresponding to the greatest confidence score (0.9826).

For each structural prediction, Boltz-2 also computes an overall confidence score on a scale from 0 to 1 that accounts for both whole-complex and interaction-interface structural confidence, with higher values indicating greater prediction confidence. Specifically, the confidence score incorporates the predicted local distance difference test (pLDDT) [11, 23] of the whole complex as well as the interface predicted template-modeling (ipTM) score [11, 24] (see Methods). Prediction confidence ranged from 0.2272 to 0.9826 across the entire set of interaction structures, with a median confidence score of 0.5516 **(Fig. 1b & c)**. Interestingly, homomeric interaction structures (*n* = 2,029; 1.86% of all structures) exhibited modestly higher confidence scores on average compared to heteromeric interaction structures (*n* = 107,253; 98.14% of all structures) **(Supplementary Fig. 1)**.

We next examined sequence length characteristics across the predicted interaction structures. Interacting protein pairs featured a broad range of sequence length combinations, with a greater overall concentration of proteins in the shorter-to-intermediate length range of approximately 200–500 amino acids **(Fig. 2a)**. Whole-complex sequence length, *i.e.*, the summed sequence length of both protein interactors, ranged from 104 to 1,998 amino acids and displayed a mild right skew, with a median of 831 amino acids **(Fig. 2b)**. To quantify differences in sequence length between interacting proteins, we calculated the sequence length ratio (*R*_SL_) according to **equation (1)**, for which values closer to 1 correspond greater similarity in interactor sequence length and values closer to 0 correspond greater sequence length disparity. Sequence length ratios were broadly distributed, with a median *R*_SL_ of 0.6072, indicating considerable variation in relative sequence length between interacting proteins **(Fig. 2c)**. In addition to sequence-level characteristics, we also considered network-level organization of the interaction structures using a focused subset of the dataset. A network graph comprising the top 1% of interaction structures ranked by confidence score revealed multiple interconnected groups formed by interactions sharing one or more protein partners **(Fig. 2d)**. This serves to illustrate the context of broader connectivity in which the binary interaction structures are situated.

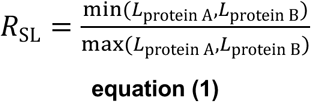

> where *R*_SL_ is the sequence length ratio,
>
> *L*_protein A_ is the sequence length of interactor protein A, and
>
> *L*_protein B_ is the sequence length of interactor protein B

### Characterization of predicted protein interaction interfaces

**Fig. 2:**
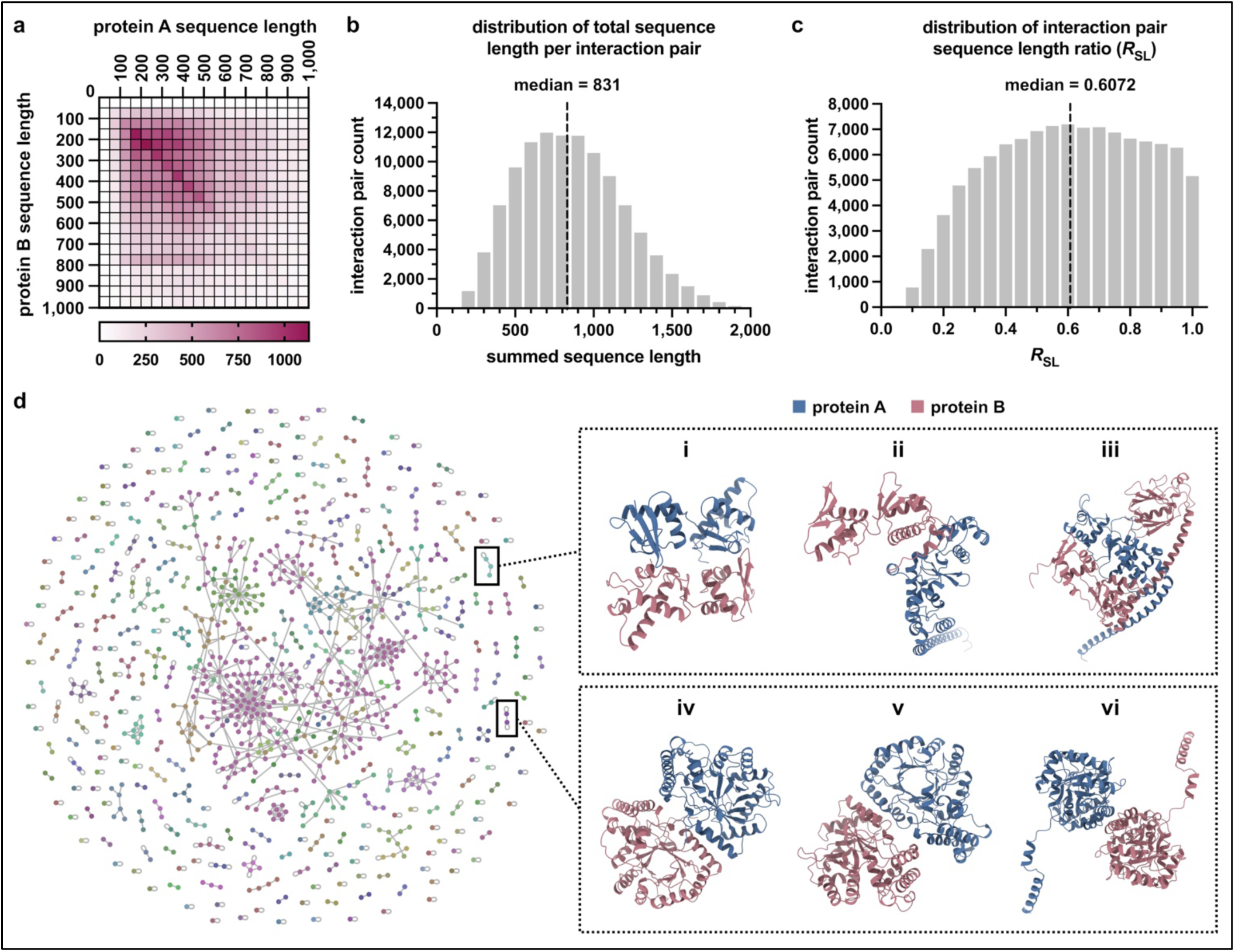
Sequence length properties of predicted protein interaction structures and examples from network organization. **a.** Distribution matrix of the sequence lengths of interacting protein pairs, with interactor protein A sequence lengths represented along the columns and interactor protein B sequence lengths along the rows. Each bin represents a 50 amino acid sequence length interval (0–1000 amino acids), and the scale bar indicates the number of predicted interaction structures. *n* = 109,282 interaction structures. **b.** Histogram of the total (summed) sequence lengths of protein A and protein B for each interaction. The median summed sequence length (831) is indicated by the dashed line. Histogram bin width = 50; *n* = 109,282 interaction structures. **c.** Histogram of sequence length ratios (*R*_SL_), calculated using **equation (1)**, across binary protein-protein interactions. *R*_SL_ values approaching 1 indicate similar sequence lengths, whereas lower values indicate greater sequence-length disparity. The median *R*_SL_ (0.6072) is indicated by the dashed line. Histogram bin width = 0.05; *n* = 109,282 interaction structures. **d.** Network representation of the top 1% (*n* = 1,093) of predicted interaction structures ranked by confidence score. Individual proteins are represented by nodes, with edges connecting interacting protein pairs. Nodes are colored according to connected-component membership, *i.e.*, proteins belonging to the same interconnected group are shown in the same color. Two interconnected groups are outlined (boxed) as examples, with the interaction structures belonging to each group depicted on the right. These include: (i) vacuolar protein-sorting-associated protein 25 (UniProt Q9BRG1) homomer; (ii) vacuolar-sorting protein SNF8 (UniPort Q96H20) with vacuolar protein-sorting-associated protein 25 (UniProt Q9BRG1); (iii) vacuolar-sorting protein SNF8 (UniProt Q96H20) with vacuolar protein-sorting-associated protein 36 (UniProt Q86VN1); (iv) aldo-keto reductase family 7 member A3 (UniProt O95154) homomer; (v) aflatoxin B1 aldehyde reductase member 2 (UniProt O43488) with aldo-keto reductase family 7 member A3 (UniProt O95154); (vi) aflatoxin B1 aldehyde reductase member 2 (UniProt O43488) homomer.

The physical interfaces of protein interactions provide the structural basis for coordinated protein association and function. To define interaction interfaces, we considered residues with a buried surface area (BSA) greater than 0 Å^2^ to be interface residues, as calculated using Proteins, Interfaces, Structures, and Assemblies (PISA) software [25, 26]. Residues meeting this criterion are reported for each interaction pair in **Supplementary Table 2**. Considering the variation in protein sequence lengths and their relative proportions within interaction pairs **(Fig. 2a-c)**, we next examined variation in their predicted interfaces. To quantify the relative extent of each interface, we calculated the interface residue fraction as the proportion of each protein’s residues located at the interaction interface, according to **equation (2) (Fig. 3a)**. Across interacting proteins, interface residue fractions showed a moderate positive correlation **(Fig. 3b)**, indicating that greater proportional interface involvement in one protein tended to be accompanied by greater involvement in its partner.

**Fig. 3:**
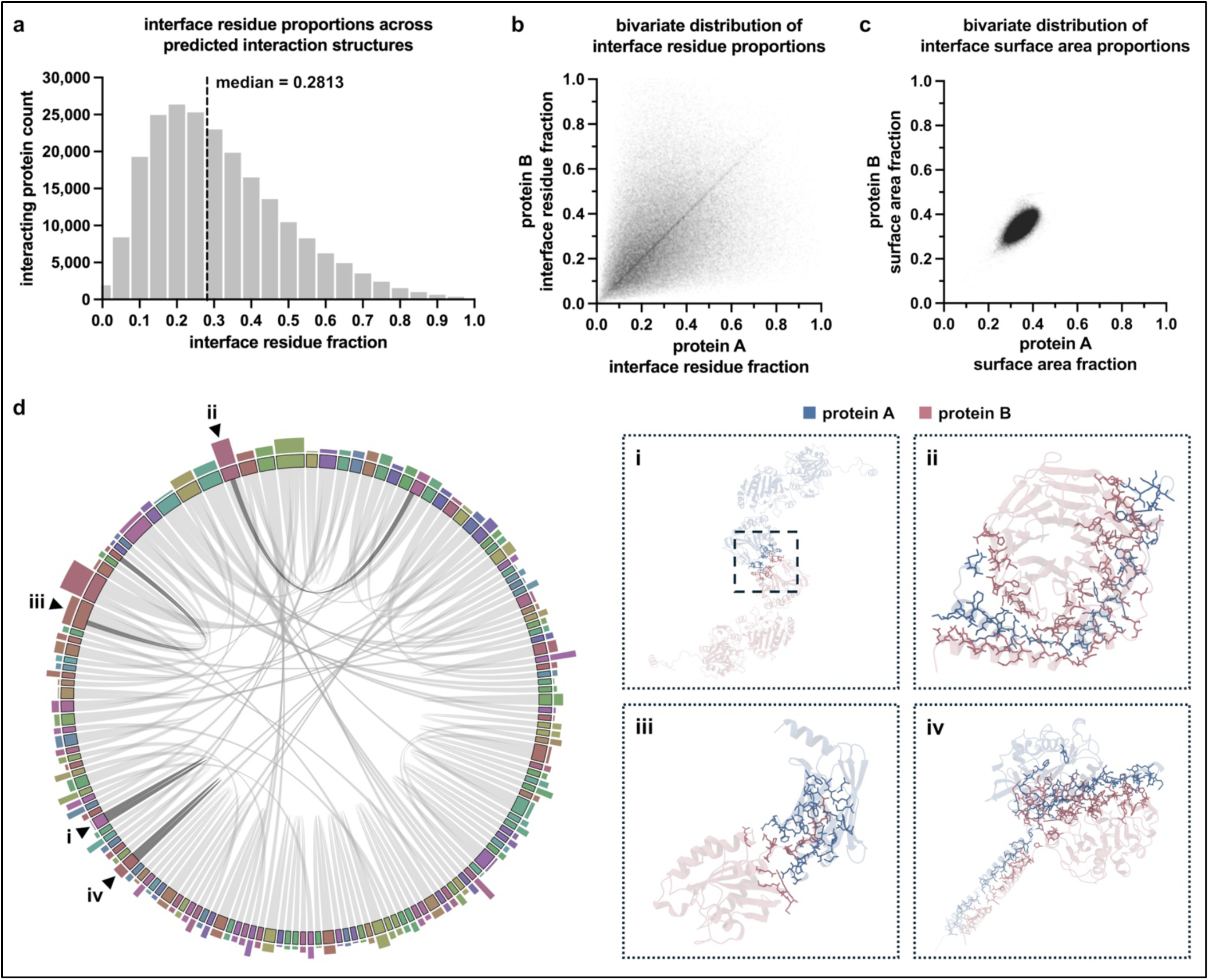
Analysis of predicted structural protein interaction interfaces. **a.** Histogram of interface residue fractions for interacting proteins across the full set of predicted interaction structures. Interface residue fractions for proteins A and B were pooled into a single distribution. The median interface residue fraction (0.2813) is indicated by the dashed line. Histogram bin width = 0.05; *n* = 218,564 interaction interface observations from 109,282 interaction structures. **b.** Bivariate distribution of interface residue fractions between the two interacting proteins in each predicted protein interaction structure. Each point represents one unique interaction. Statistical analysis by Spearman correlation (*r*_S_ = 0.4650, 95% CI 0.4602 to 0.4698, *p* < 0.0001) and Pearson correlation (*r* = 0.4077, 95% CI 0.4028 to 0.4127, *p* < 0.0001), *n* = 109,282 interaction structures. **c.** Bivariate distribution of interface surface area fractions between the two interacting proteins in each predicted protein interaction structure. Each point represents one unique interaction. Statistical analysis by Spearman correlation (*r*_S_ = 0.6679, 95% CI 0.6645 to 0.6713, *p* < 0.0001) and Pearson correlation (*r* = 0.7985, 95% CI 0.7963 to 0.8006, *p* < 0.0001), *n* = 109,282 interaction structures. **d.** Chord diagram of the top 0.01% (*n* = 109) of predicted interaction structures ranked by confidence score. Interactor proteins are represented by sectors along the plot circumference, with chords connecting interacting protein pairs. Outer bar heights correspond to the mean interface residue fraction of each protein, normalized to the greatest mean in the plot. Chords of example interaction structures are depicted in dark grey, with the respective structures and atom representations of interface residues shown to the right. These include structures with: (i) the minimum interface residue fraction, 0.0285 (interface residues outlined by dashed box), showing procollagen-lysine,2-oxoglutarate 5-dioxygenase 2 (UniProt O00469) homomer; (ii) the maximum interface residue fraction, 0.7826, showing guanine nucleotide-binding protein subunit gamma-T2 (UniProt O14610) in complex with guanine nucleotide-binding protein subunit beta-2 (UniProt P62879); (iii) the lower median-adjacent interface residue fraction, 0.2800, showing retinal rod rhodopsin-sensitive cGMP 3’,5’-cyclic phosphodiesterase subunit delta (UniProt O43924) in complex with Ras-related protein Rap-2b (UniProt P61225); and (iv) the upper median-adjacent interface residue fraction, 0.2866, showing galactosylgalactosylxylosylprotein 3-beta-glucuronosyltransferase 3 (UniProt O94766) homomer, all relative to the median interface residue fraction of 0.2813 from panel **a**.

In a similar manner, interface surface area fraction was calculated as the proportion of BSA relative to the combined BSA and ASA for each protein in a given interaction, according to **equation (3)**. A strong positive correlation was observed between the interface surface area fractions of interacting proteins, with values concentrated within a relatively narrow range for both partners **(Fig. 3c)**, conveying a tendency toward coordinated proportional surface burial across interaction partners. To illustrate the variation in interface extent across a subset of interactions, the top 0.01% of interaction structures ranked by prediction confidence were visualized through a chord diagram **(Fig. 3d)**. This representation highlights the heterogeneity of the predicted interaction interfaces, with different interactions displaying varying degrees of residue involvement. These analyses delineate interface-localized residues across the predicted interaction structures and provide insight into the relative extent of their interfaces.

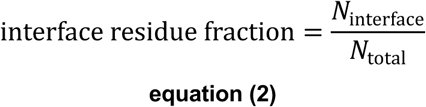

> where *N*_interface_ is the number of interface residues,
>
> *N*_total_ is the total number of residues in the protein, and
>
> the interface residue fraction is calculated separately for proteins A and B

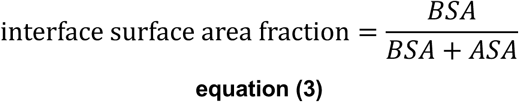

> where *BSA* is the buried surface area of a protein in Å^2^,
>
> *ASA* is the accessible surface area of a protein in Å^2^, and
>
> the surface area fraction is calculated separately for proteins A and B
>
> (with the sum of BSA and ASA representing the total surface area of a given protein)

We further characterized the predicted interaction interfaces by examining intermolecular bonding, *i.e.* between proteins, across the interaction structures using PISA **(Supplementary Table 2)**. Hydrogen bonds and salt bridges were quantified as bond densities relative to mean interface BSA, as outlined in **equation (4)**. Hydrogen bond and salt bridge densities exhibited medians of 6.634 and 3.127 bonds per 1,000 Å², respectively, with salt bridge densities observed more frequently at lower values than hydrogen bond densities across distributions **(Fig. 4a-c)**. The larger proportion of interaction structures with greater hydrogen bond densities is consistent with the broader range of functional groups capable of hydrogen bond formation, whereas salt bridges depend on the proximity and orientation of oppositely charged groups at the interaction interface. In addition to intermolecular bonding, we characterized the energetic properties of predicted interaction interfaces through PISA-based estimation of solvation and stabilization energies **(Supplementary Table 2)**. Both energetic properties exhibited a left-skewed, predominantly negative distribution, with median values of -34.81 and -52.23 kcal/mol for solvation and stabilization, respectively **(Fig. 4d)**. These distributions suggest that favorable solvation and stabilization contributions were common among the predicted interaction structures, and may partly reflect the experimental origin of the underlying interaction pairings.

**Fig. 4:**
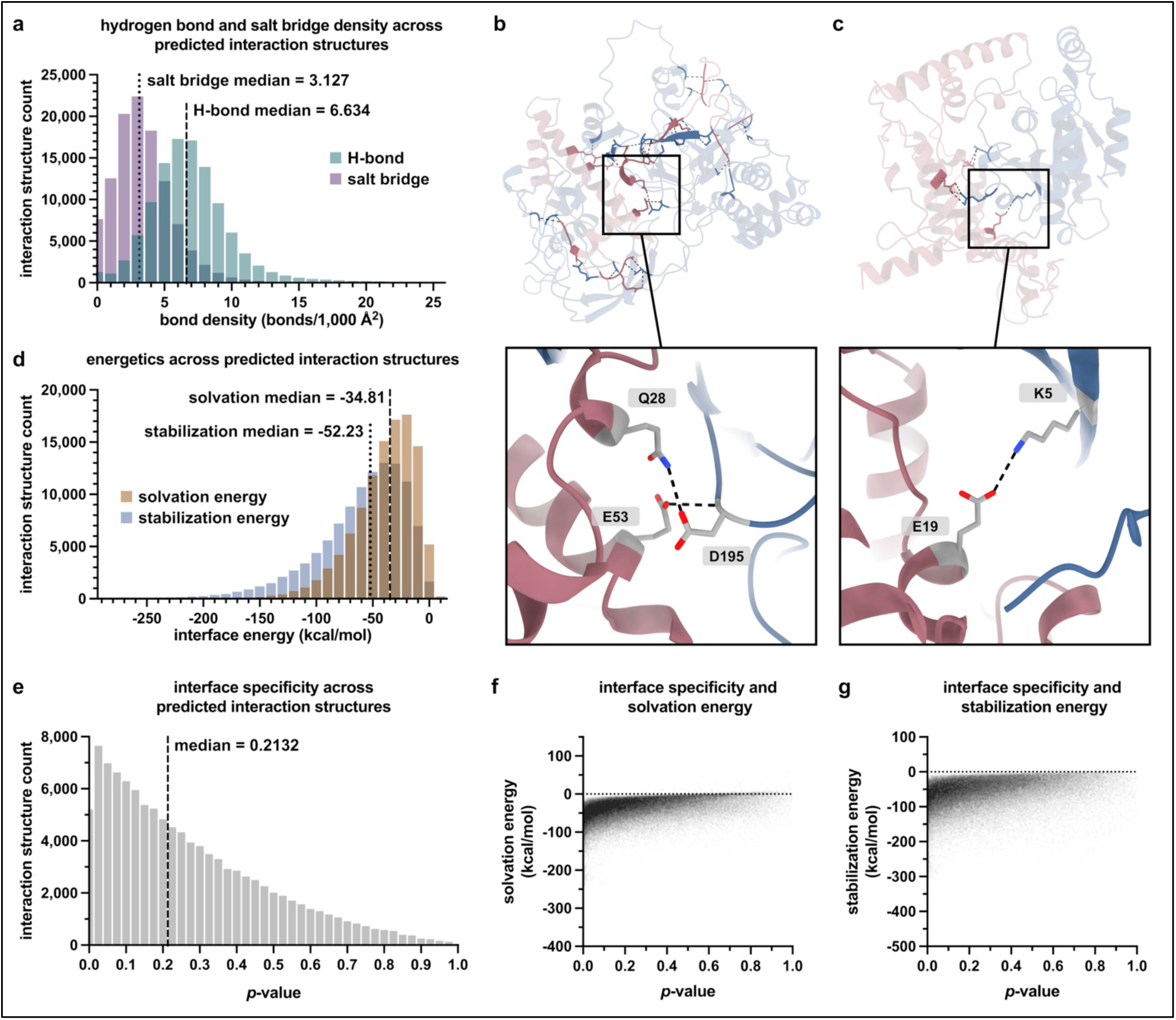
Intermolecular bonding and energetic properties. **a.** Histogram of intermolecular hydrogen bond and salt bridge densities across predicted interaction structures. Hydrogen bond and salt bridge densities are displayed as separate distributions, distinguished by color. Bond densities are quantified as bonds per 1,000 Å² of mean BSA, as specified in **equation 4**. Medians are indicated by the dashed and dotted lines for hydrogen bond density (6.634) and salt bridge density (3.127), respectively. Histogram bin width = 1.0; *n* = 109,282 interaction structures. The upper limit of the *x*-axis was set to the 99.9^th^ percentile of the hydrogen bond density distribution for visual interpretability of the predominant range of bond densities; all data points, including those above this threshold, were retained for statistical analysis. **b-c.** Representative structures corresponding to the lower median-adjacent bond density values for: (b) hydrogen bond density, histone-lysine N-methyltransferase KMT5C (UniProt Q86Y97) in complex with histone H4 (UniProt P62805); and (c) salt bridge density, cell division control protein 42 homolog (UniProt P60953) in complex with zinc finger protein 420 (UniProt Q8TAQ5). Part of the zinc finger protein 420 protein that is distal to the interaction interface is not shown for visual clarity. **d.** Histogram of solvation and stabilization energies across predicted interaction structures, displayed as separate distributions distinguished by color. Medians are indicated by the dashed and dotted lines for solvation energy (-34.81) and stabilization energy (-52.23), respectively. Histogram bin width = 10; *n* = 109,282 interaction structures. The *x*-axis was restricted to the range between the lower of the two distributions’ 0.01^st^ percentile values and the higher of their 99.9^th^ percentile values for visual interpretability of the predominant ranges; all data points, including those above this threshold, were retained for statistical analysis. **e.** Histogram of the PISA-calculated *p*-values across interaction structures. The median p-value (0.2132) is indicated by the dashed line. Histogram bin width = 0.025; *n* = 108,593 interaction structures. **f-g**. Relationships between *p*-value and **(f)** solvation energy or **(g)** stabilization energy. Each point represents one unique interaction. Statistical analysis by Spearman correlation (solvation: *r*_S_ = 0.4695, 95% CI 0.4647 to 0.4743, *p* < 0.0001; stabilization: *r*_S_ = 0.3446, 95% CI 0.3392 to 0.3500, p < 0.0001) and Pearson correlation (solvation: *r* = 0.3974, CI 0.3924 to 0.4024, *p* < 0.0001; stabilization: *r* = 0.2783, CI 0.2728 to 0.2838, *p* < 0.0001), *n* = 108,593 interaction structures. For *n* = 689 interaction structures, *p*-values were undetermined and therefore not included in the plots in panels **e-g** (see Methods).

We also considered the PISA *p*-value, a statistical measure of interface specificity, representing the probability of obtaining a lower solvation energy if interface atoms were instead randomly selected from the protein surface [25]. A *p*-value below 0.5 indicates greater-than-expected interface hydrophobicity and is considered to indicate that the interface is interaction-specific [25]. 91,730 (83.9%) of the predicted interaction structures had a *p*-value less than 0.5, with the median *p*-value at 0.2132 **(Fig. 4e)**. Additionally, *p*-values exhibited a moderate positive correlation with solvation energy and a weaker, modest positive correlation with stabilization energy, indicating a tendency for lower *p*-values to accompany more negative energetic estimates. These associations are consistent with the interrelatedness of *p*-value, solvation energy, and stabilization energy, while the metrics themselves represent distinct but complementary properties of the predicted interaction interfaces. Overall, these analyses characterize the structural extent, intermolecular bonding, energetic properties, and hydrophobic specificity of the predicted interaction interfaces.

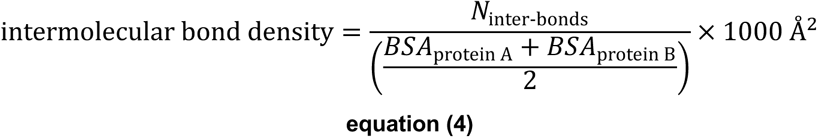

> where *N*_inter-bonds_ is the number of intermolecular hydrogen bonds or salt bridges (calculated independently),
>
> *BSA*_protein A_ is the buried surface area of interactor protein A in Å^2^,
>
> *BSA*_protein B_ is the buried surface area of interactor protein B in Å^2^, and
>
> the density is considered per 1000 Å^2^ of surface area

### Structural mapping of pathogenic mutations

The localization of pathogenic mutations within protein interaction interfaces provides a structural perspective on their potential involvement in disease-associated protein interactions. To investigate this across the predicted interaction structures, we mapped clinically pathogenic missense variants sourced from ClinVar [22, 27] to the interaction interfaces, identifying interface-localized mutations associated with 4,306 disease categories **(Supplementary Tables 3 & 4)**. It should be noted that these disease categories directly correspond to distinct ClinVar disease annotations but may include overlapping or related clinical terminology, yet original annotations were preserved to maintain consistency with ClinVar data. The mapping of interface-localized mutations was further explored within the disease contexts of acute myeloid leukemia (AML), Alzheimer’s disease (AD), and cardiomyopathy (CM), focusing on interactions within the upper quartile of ipTM scores, the interface-specific component of the Boltz-2 confidence score (see Methods). The distribution of interface mutation-involving interactions within these disease contexts revealed highly connected proteins, which we refer to as "hub proteins" **(Fig. 5a)**. Hub proteins identified include CCAAT/enhancer-binding protein alpha (C/EBPα) in AML, amyloid-β precursor protein (APP) in AD, and phospholamban in CM.

**Fig. 5:**
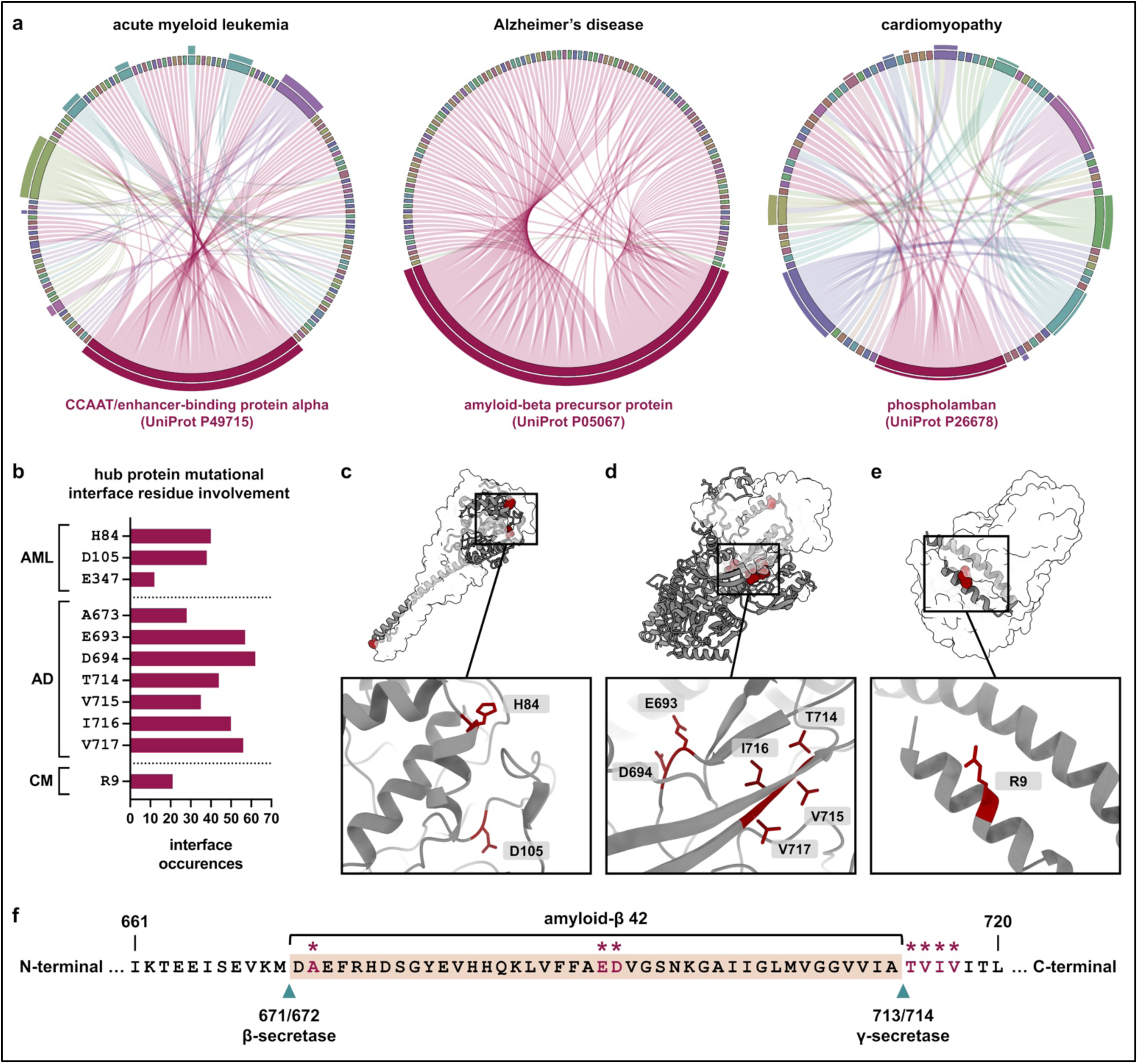
Interface-localized mutations within disease contexts. **a.** Chord diagrams of interactions involving interface-localized pathogenic mutations associated with acute myeloid leukemia (AML), Alzheimer’s disease (AD), and cardiomyopathy (CM), restricted to interactions with ipTM scores in the upper quartile of the entire dataset (above Q3). Sectors represent individual proteins, and chords connect interacting pairs, with each chord colored to match the sector of the mutation-bearing protein. The sectors of hub proteins, *i.e.*, proteins involved in the greatest number of interactions, are annotated and colored magenta. Outer bar heights correspond to the mean number of interface mutational residues per interaction involving each protein, normalized to the greatest mean for each plot. **b.** The number of occurrences of hub protein interface mutational residues across interactions represented in panel **a** for AML, AD, and CM. **c-d.** Representative interaction structures containing all interface mutational residues for each pathology, including: **(c)** AML, hub protein CCAAT/enhancer-binding protein alpha (UniProt P49715) in complex with DNA damage-inducible transcript 3 protein (UniProt P35638); **(d)** AD, hub protein amyloid-beta precursor protein (UniProt P05067) in complex with Cyclin-G1 (UniProt P51959); and **(e)** CM, hub protein phospholamban (UniProt P26678) in complex with Bcl-2-like protein 13 (UniProt Q9BXK5). Hub proteins are shown in grey and interactor proteins are semi-transparent for visualization of interface mutational residues, which are depicted in red. **f.** Amino acid sequence of the APP region containing the Aβ42 peptide, highlighted in beige (UniProt P05067). Arrowheads indicate cleavage sites of β-secretase and γ-secretase. The seven interface-localized mutational residues of APP are denoted by asterisks.

C/EBPα is a transcription factor that regulates cellular differentiation and metabolic processes while also suppressing cellular proliferation [28, 29], with its dysregulation known to contribute to the development of AML [30-32]. APP is a multifunctional protein involved in cellular adhesion and signalling, with important roles in nervous system development, synaptic function, as well as neuronal survival [33]. The proteolytic processing of APP generates amyloid-β peptides, including the aggregation-prone amyloid-β 42 peptide (Aβ42) which is implicated in AD pathogenesis [34-36]. Phospholamban modulates cardiac relaxation and contractility by regulating sarcoplasmic reticulum calcium uptake [37], with pathogenic variants implicated in various forms of CM [38-40]. Interface-localized mutational residues were found among multiple interactions involving these hub proteins, exhibiting varying frequencies of occurrence across interactions **(Fig. 5b-e)**. APP exhibited the greatest diversity of interface-localized mutational residues, followed by C/EBPα, whereas phospholamban was represented by a single mutational residue across its associated interactions. Although these distributions may be biologically meaningful, they should be interpreted with consideration of database annotation bias, given that BioGRID [6] and ClinVar [22] depend on the availability of previously characterized protein interactions and pathogenic variants, respectively.

Of particular interest, the interface-localized mutational residues of APP were concentrated within and immediately adjacent to the AD-associated Aβ42 region, which spans residues 672-713 [36] **(Fig. 5f)**. Production of Aβ42 involves cleavage of APP by β-secretase at M671/D672 [36, 41, 42] followed by sequential γ-secretase processing at multiple sites, culminating in cleavage at A713/T714 to generate the Aβ42 C-terminus [36, 43, 44]. The relative positions of the interface-localized mutational residues include: A673 in close proximity to the β-secretase cleavage site; E693 and D694 centrally located within Aβ42; and T714, V715, I716, and V717 adjacent and proximal to the γ-secretase cleavage site **(Fig. 5f)**. This positioning highlights a notable correspondence between predicted interaction interfaces and regions important for APP processing with potential relevance to AD pathogenesis. Collectively, these disease-focused analyses provide predicted structural contextualization of pathogenic variation across human protein interactions which may be useful for the study and interpretation of disease mechanisms.

### Confidence, experimental structure comparisons, limitations, and opportunities for learning

Confidence metrics provide useful information for evaluating the quality of predicted protein interaction structures. Boltz-2 provides an overall confidence score alongside pTM and pLDDT, which reflect global and local structural confidence, respectively, and their interface-specific counterparts, ipTM and ipLDDT. The distributions of these metrics are shown in **Fig. 6a** and reported in **Supplementary Table 5**. Overall, the metrics exhibited considerable variability, with higher median values for pLDDT and ipLDDT than for pTM and ipTM. The particularly low ipTM values suggest uncertainty in the predicted inter-chain topology, *i.e.*, the relative arrangement of subunits. Previous studies suggest that ipTM scores are sensitive to several factors, including including disordered regions, non-interacting domains, transient protein interactions, and incomplete representation of higher-order molecular assemblies [45-47], each of which may partly contribute to the relatively low ipTM values observed here.

**Fig. 6:**
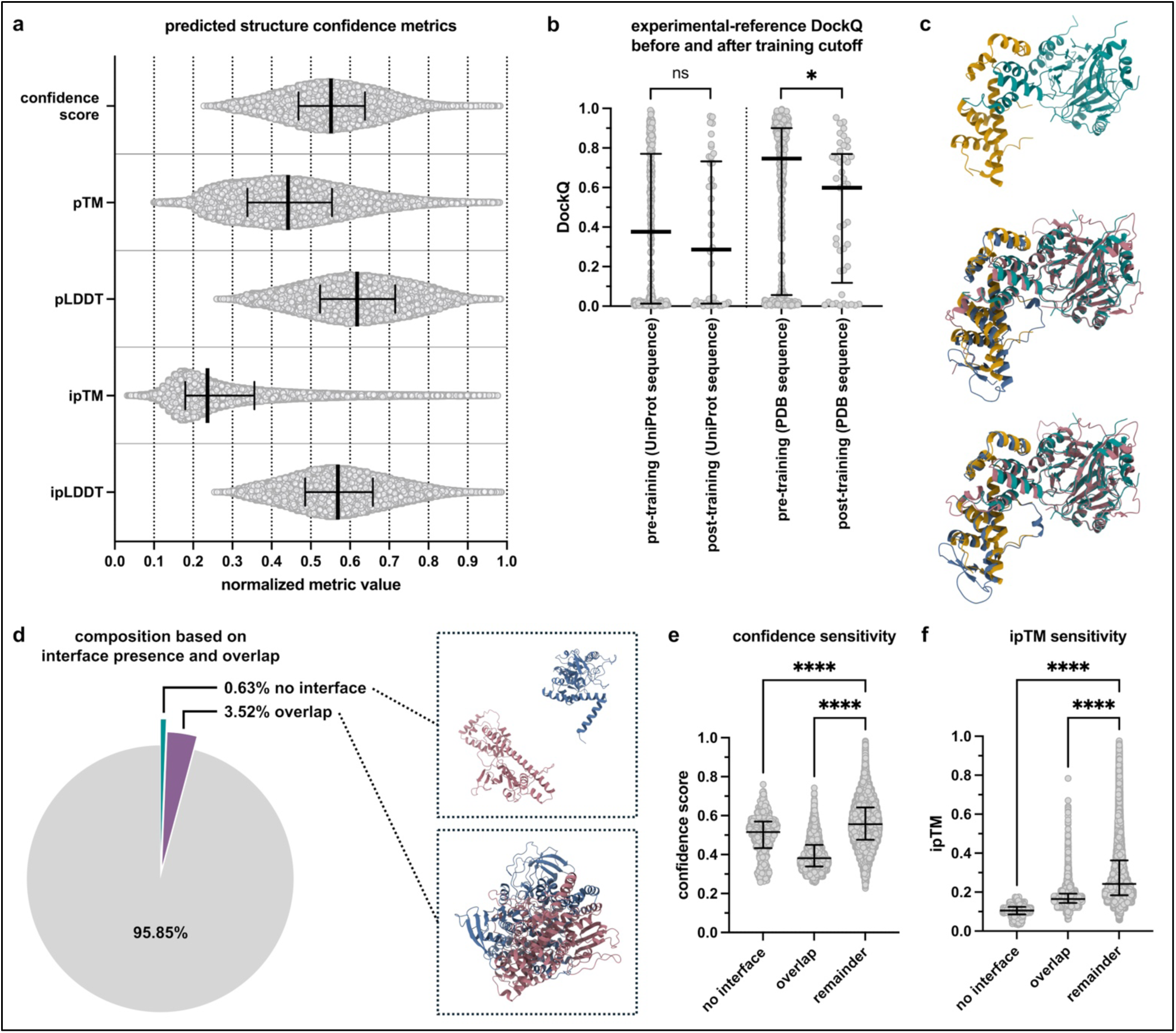
Prediction confidence, experimental comparisons, and abnormal structures. **a.** Distribution of confidence metrics for the *n* = 109,282 predicted interaction structures. Each point represents an individual interaction structure, with medians and IQRs shown. **b.** DockQ scores comparing experimental PDB structures to corresponding interaction structure predictions based on UniProt sequence from the main prediction dataset (grey) and to interaction structure predictions based on the exact sequences of the PDB structures (magenta). DockQ comparisons are separated into pre- and post-training groups based on whether the PDB structure predates or postdates the Boltz-2 training cutoff (see Methods for further details). Groupings include: *n* = 281 pre-training (UniProt sequence); *n* = 42 post-training (UniProt sequence); *n* = 370 pre-training (PDB sequence); and *n* = 45 post-training (PDB sequence). For UniProt sequence-based interaction structures with more than one corresponding PDB structure, the mean DockQ was used, resulting in fewer datapoints for UniProt sequence than for PDB sequence DockQ. Data represented as medians and IQRs. Statistical analysis by Mann-Whitney test, \**p* = 0.0122, ns, not significant. **c.** Example of a post-training interaction structure of S-phase kinase-associated protein 1 (UniProt P63208) in complex with F-box only protein 22 (UniProt Q8NEZ5). The experimentally-determined structure (PDB 8S7E) of this interaction is shown with S-phase kinase-associated protein 1 depicted in yellow and F-box only protein 22 depicted in teal (top). DockQ = 0.7247 between the experimental structure and UniProt-sequence based prediction (middle), while DockQ = 0.8049 between the experimental structure and PDB-sequence based prediction (bottom). For both predicted structures, S-phase kinase-associated protein 1 is depicted in blue and F-box only protein 22 is depicted in red. **d.** Composition of the predicted interaction structure dataset (*n* = 109,282) depicting amount of structures with no detected interface in teal (*n* = 689 structures; 0.63%) and amount of structures with structural overlap in purple (*n* = 3,846 structures; 3.52%), defined as 10% or more of either interacting protein having inter-protein steric clashes (see methods). Example predicted structures are shown for 1-acyl-sn-glycerol-3-phosphate acyltransferase beta (UniProt O15120) in complex with PHD finger protein 11 (UniProt Q9UIL8), which had no detectable interface (top box), and coilin (UniProt P38432) in complex with testis-specific Y-encoded-like protein 2 (UniProt Q9H2G4), which exhibited structural overlap (bottom box). **(e-f)** Comparisons of **(e)** confidence scores and **(f)** ipTM among interaction structures with no detectable interface (*n* = 689), structural overlap (*n* = 3,846), or neither condition (remainder; *n* = 104,747). Data are represented as individual interaction structures with medians and IQRs. Statistical analysis by Kruskal-Wallis test followed by Dunn’s multiple comparisons test; \*\*\*\**p* < 0.0001.

To further evaluate the predicted interaction structures, we performed comparisons against experimentally determined structures obtained from the Protein Data Bank (PDB) [48, 49], specifically binary interaction structures to maintain a consistent structural context with the binary predictions. Structural agreement was quantified by DockQ [50] and experimental structures separated according to whether they were released before or after the Boltz-2 training cutoff of June 1, 2023 [14]. Comparisons were conducted separately for predictions based on UniProt sequences (from the main dataset) and predictions based on the corresponding experimentally determined PDB structure sequences **(Fig. 6b & c; Supplementary Tables 6 & 7)**, considering that variations in sequence composition of experimental structures may influence structural agreement. Median DockQ scores were lower in the post-training groups for both UniProt sequence- and PDB sequence-based predictions, although this difference was statistically significant only for PDB sequence-based predictions, which also exhibited higher median DockQ scores than UniProt sequence-based predictions in both the pre- and post-training groups. These comparisons provide insight into prediction accuracy, although differences in sequence composition may limit the direct comparability of predicted and available experimental structures, a factor that should be considered when interpreting the extent of structural agreement.

Finally, we examined the prevalence of pronounced structural abnormalities, namely the absence of interaction interfaces and aberrant inter-protein structural overlap. Among the entire set of 109,282 interaction structures, 689 (0.63%) lacked a detectable interface, while 3,846 (3.52%) exhibited structural overlap involving at least 10% of residues in either interacting protein **(Supplementary Table 8; Fig. 6d)**. Notably, overall confidence scores were particularly sensitive to structural overlap, whereas ipTM values were particularly sensitive to the absence of detectable interaction interfaces **(Fig. 6e & f)**. These subsets of structurally abnormal predictions may serve as informative training data for future machine learning applications aimed at refining structural confidence assessments and improving interaction structure prediction accuracy. Together, these analyses provide insight into the overall quality and limitations of the predicted interaction structures, highlighting the value of complementary assessments for evaluating structural reliability and how they may inform future computational approaches. Regardless of prediction quality and structural abnormalities, all interaction structures were retained throughout the analyses performed in this study.

## Discussion

The large-scale structural modelling and analysis presented here provide structural contextualization of experimentally supported human protein interactions and integrate this information with disease-associated mutation data. Reflecting the diversity of the human proteome, the predicted interaction structures exhibited considerable heterogeneity in terms of interface composition, intermolecular bonding, and energetic properties. The structural mapping of pathogenic variants localized mutational residues to predicted protein interaction interfaces across disease categories and revealed highly connected proteins within the exemplified disease contexts. Assessments of prediction confidence, agreement with experimentally determined structures, and the presence of structural abnormalities indicated variable predictive reliability, emphasizing the importance of accounting for uncertainty when interpreting predictions. It should be noted that protein interaction structures can be influenced by various biological factors, such as phosphorylation, other post-translational modifications, and the broader cellular environment, the effects of which are not explicitly captured by the modelling performed here and extend beyond the scope of the current work. Despite limitations, these findings point to opportunities offered by large-scale structural modelling in the study of the human protein interactome.

The broad range of protein sequence lengths and relative interactor sizes represented among the predicted structures reflects the intra- and inter-molecular diversity of human protein interactions [2, 3]. Interestingly, positive correlations between the interface proportions of interacting partners, particularly for proportional surface burial, suggest a tendency toward corresponding relative interface involvement, despite differences in the size and composition of interactors. The predominance of hydrogen bonds relative to salt bridges, together with generally favorable energetic estimates and interface specificity measures, suggests that many predicted interfaces exhibit physicochemical features consistent with experimental observation [25]. These characterizations provide insight into the commonalities and differences underlying the predicted structural organization of human protein interactions.

The mapping of pathogenic amino acid variants to the interaction interfaces provides predicted structural insight into the potential significance of disease-associated mutations. ClinVar is a particularly valuable resource for this purpose as it provides clinically grounded data with extensive variant coverage [21, 22], extending the potential utility of structural interaction modelling to translational research. Within the disease contexts of AML and CM, the distributions of interface mutation-involving interactions extended beyond involvement of the C/EBPα and phospholamban hub proteins, respectively, indicating greater diversity in interactions that are potentially relevant to these diseases. In contrast, APP emerged as a nearly universal hub in AD, accounting for almost all interface mutation-involving interactions. In relation to AD pathogenesis, the localization of APP interface mutations within and adjacent to the Aβ42 region identifies a potentially meaningful structural association that may inform investigations into amyloidogenic processing.

With respect to prediction quality, the various factors considered in this study provide complementary perspectives on structural reliability. The variability in confidence metrics across the dataset warrants caution against interpreting the predicted interaction structures as equally reliable. Comparisons with experimentally determined structures yielded mixed results, with differences between predicted and experimental structure sequence composition potentially contributing to these outcomes and complicating the assessment of prediction accuracy. Furthermore, the occurrence of structural abnormalities, *i.e.*, interface absence and structural overlap, identifies cases in which computational predictions fail to produce plausible interaction structures, despite underlying experimental support for the interacting protein pairs. Given the systematic scope of this work, predicted interaction structures with these abnormalities were not excluded from analyses presented in the main text (see Methods). Analyses excluding these structures exhibited broadly consistent results without major deviations in distributions and trends, including for interface proportions **(Supplementary Fig. 2)**, bonding and energetic properties **(Supplementary Fig. 3)**, and interface specificities **(Supplementary Fig. 4)**. The structural abnormalities observed may also inform future development of AI/ML-based approaches for interaction structure prediction, particularly through the use of structurally implausible predictions as negative training examples. Additionally, the incorporation of experimentally supported interaction pair data into model training may represent a complementary avenue for improving prediction accuracy.

In summary, the structural modelling and analyses presented here provide a large-scale resource for investigating experimentally supported human protein interactions at the molecular level. The systematic characterization of predicted interaction interfaces, together with the structural mapping of pathogenic mutations, offers a basis for investigating relationships between protein interaction structure, function, and disease. Although prediction uncertainties necessitate cautious interpretation, these findings provide a foundation for further hypothesis generation and experimental investigation. As computational prediction methods continue to improve, the integration of large-scale structural modelling with experimental approaches are anticipated to enable increasingly detailed research on the human protein interactome and its involvement in disease.

## Methods

### Integration of Protein Interaction and Sequence Data

Protein interaction data were obtained from the BioGRID database file repository [6, 51] in Molecular Interaction TAB (MITAB) tab-delimited format [52]. The original ordering of interaction partners as protein A and protein B was maintained for consistency. A number of filtering criteria were applied to the BioGRID data to retain interactions consistent with the scope of the study and suitable for subsequent structural predictions. These criteria included: (i) inclusion of only interactions classified as “direct interaction” (PSI-MI term MI:0407), which is reserved for interactions between molecules that are in direct contact with each other [52, 53]; (ii) inclusion of only interactions for which both interacting proteins originate from *Homo sapiens*, and (iii) exclusion of redundant interactions, with reciprocal representations of the same protein pair also treated as redundant, to retain a single instance of each unique interaction. Amino acid sequences corresponding to each protein were retrieved using the UniProt website REST API [19] by UniProt identifiers. Interactions for which either protein sequence contained selenocysteine (U) or pyrrolysine (O) were excluded such that all retained sequences were composed exclusively of the 20 standard amino acids, as required for structural prediction. This processing resulted in a non-redundant dataset of direct human protein interaction pairs and their amino acid sequences, comprising 109,282 binary interactions.

### Structural Prediction of Protein Interactions

Three-dimensional structures of the binary protein interactions were predicted using Boltz-2 [14] version 2.2.1. Computations were performed in Google Colab using the High-RAM configuration together with an A100 GPU for hardware acceleration. For each interaction, the amino acid sequences of the two interacting proteins were provided as input to Boltz-2 for joint complex prediction as separate protein chains (designated A and B), preserving the protein A/protein B interaction-partner ordering retained during BioGRID processing. Default Boltz-2 parameters were used, including one diffusion sample, three recycling steps, 200 sampling steps, and a diffusion step scale of 1.5. No structural templates or additional structural constraints were provided as prediction inputs. MSA generation was enabled with the default greedy MSA pairing strategy applied. MSA generation utilized the ColabFold MSA server (https://api.colabfold.com), which employs MMseqs2 for sequence homology search and MSA construction [54, 55]. Confidence metrics were computed by Boltz-2 for each predicted complex, with the overall confidence score defined as follows: confidence score = 0.8(complex pLDDT) + 0.2(ipTM) [14]. The above prediction configurations were applied consistently across the entire interaction dataset, and the resulting interaction structures were retained in mmCIF format for subsequent structural analyses.

### Interaction Network, Structural Interface, and Mutational Analyses

The protein interaction network in **Fig. 2d** was constructed using the igraph Python package [56], with individual proteins represented as nodes and binary interactions represented as edges. Network visualization was performed using the graphopt layout algorithm. Structural interfaces were systematically characterized using PISA software [25]. Interface residues were defined as those for which BSA was greater than 0 Å^2^, as determined by PISA. Interface surface area fractions and intermolecular bond densities were calculated using values determined by PISA, as outlined in **equations (3) and (4)**, respectively. For 689 structures (0.63% of all structures), interfaces were not detected by PISA and *p*-values were therefore undetermined. Consequently, these 689 structures were excluded from the *p*-value plots in **Fig. 4e-g**. The solvation and stabilization energies for these 689 structures were assigned values of zero, equating the absence of a detectable interface with negligible energetic contributions, and were included in the distribution plot in **Fig. 4d**. Chord diagrams were generated using the pyCirclize Python package [57], with individual proteins represented as sectors spanning the plot circumference and chords representing interaction-associated interface or mutational features, as indicated for each analysis.

Mutational variants were retrieved according to UniProt identifier using the EMBL-EBI Proteins API [58] and cross-referenced with variant-disease pair Reference ClinVar (RCV) records obtained from the ClinVar [22] NCBI website. Analysis was restricted to missense mutational variants classified as “Pathogenic” and having an associated disease annotation. Mutational amino acid residues were mapped onto their corresponding predicted interaction structures with localization at the interaction interface based on the PISA-computed residue BSA > 0 Å^2^ criteria described above. Identified interface variants were subsequently grouped according to their associated disease annotations as designated in the RCV record obtained from ClinVar.

### Comparison to Experimental Structures and Identification of Interface Abnormalities

Experimental interaction structures from the RCSB PDB [48, 49] used as reference structures in evaluating the predicted interaction structures were identified by UniProt identifiers using the RCSB PDB Search API [48]. Experimental structures were filtered to retain only unmutated, two-chain protein complexes with standard amino-acid sequences and were divided according to whether they were released before (pre-training) or after (post-training) the Boltz-2 training cutoff of June 1, 2023 [14]. Two separate comparisons against the PDB structures were performed: one using interaction structures predicted from the UniProt sequences of the main dataset and the other using interaction structures predicted from the sequences of the PDB entries themselves. Structural similarity between experimental and predicted interaction structures was evaluated using DockQ v2 [50]. Where multiple PDB structures were available for the same interaction from the main (UniProt sequence-based) dataset, DockQ scores were averaged to obtain a single value for that interaction, and DockQ scores were compared with Boltz-2 confidence metrics.

Abnormalities that were evaluated include the absence of structural interfaces and inter-protein structural overlap. Interaction structures were classified as lacking interfaces when no interface was detected between the two protein chains by PISA [25]. Structural overlap was evaluated using the Biopython Bio.PDB package [59]. Inter-protein steric clashes were defined as atomic contacts occurring at distances ≤ 50% of the combined van der Waals radii of the two atoms, as obtained from Bickerton et al. [60]. For each protein, the percentage of residues involved in at least one inter-protein steric clash was calculated relative to its total residue count. Interaction structures were classified as exhibiting substantial structural overlap when ≥ 10% of residues in either interacting protein were involved in steric clashes. Regardless of prediction quality and structural abnormalities, all interaction structures were retained throughout the various analyses performed in alignment with the systematic scope of this work unless otherwise stated in the Figure legends.

### Statistical Analyses

Statistical analyses and graphical representations were performed using GraphPad Prism 11. These analyses included and Spearman and Pearson correlations, Mann-Whitney tests, Kruskal-Wallis tests, and descriptive statistics, with various data distributions visualized using histograms. Details of the statistical analyses of individual datasets are provided in the corresponding figure legends.

## Data Availability

The predicted interaction structures generated in this study are available online through FigShare: https://doi.org/10.6084/m9.figshare.34842945

## Acknowledgments

This work was supported in part by funds provided through the Canada Impact+ Research Training Awards (CIRTA) Program issued to A.M.I by the Natural Sciences and Engineering Research Council of Canada (NSERC). We gratefully acknowledge the individuals and organizations responsible for developing and maintaining BioGRID [6, 9], the UniProt Knowledgebase [19, 61], the ClinVar database [21, 22], and the RCSB PDB [48, 49] as valuable, freely accessible resources for the scientific community. We also thank the developers of Boltz-2 [14] and PISA [25, 26], both of which were instrumental to conducting this research. Molecular visualizations were generated with UCSF ChimeraX [62], developed by the RBVI at the University of California, San Francisco.

