## Supplemental Information Document for "AI-based structural modelling and pathogenic variant mapping across the human protein interactome"

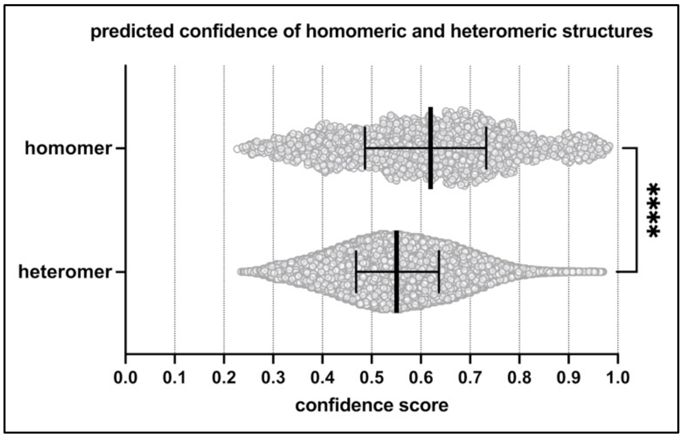


**Supplementary Fig. 1: Homomeric and Heteromeric prediction confidence.** Distribution of confidence scores across predicted protein interaction structures for homomers and heteromers. Each point represents one interaction structure. Data are represented as medians of confidence scores (homomer median: 0.6199; heteromer median: 0.5506) and interquartile ranges. Statistical analysis by Mann-Whitney test, ****$p < 0.0001$. $n = 2{,}029$ homomeric interaction structures and $n = 107{,}253$ heteromeric interaction structures.

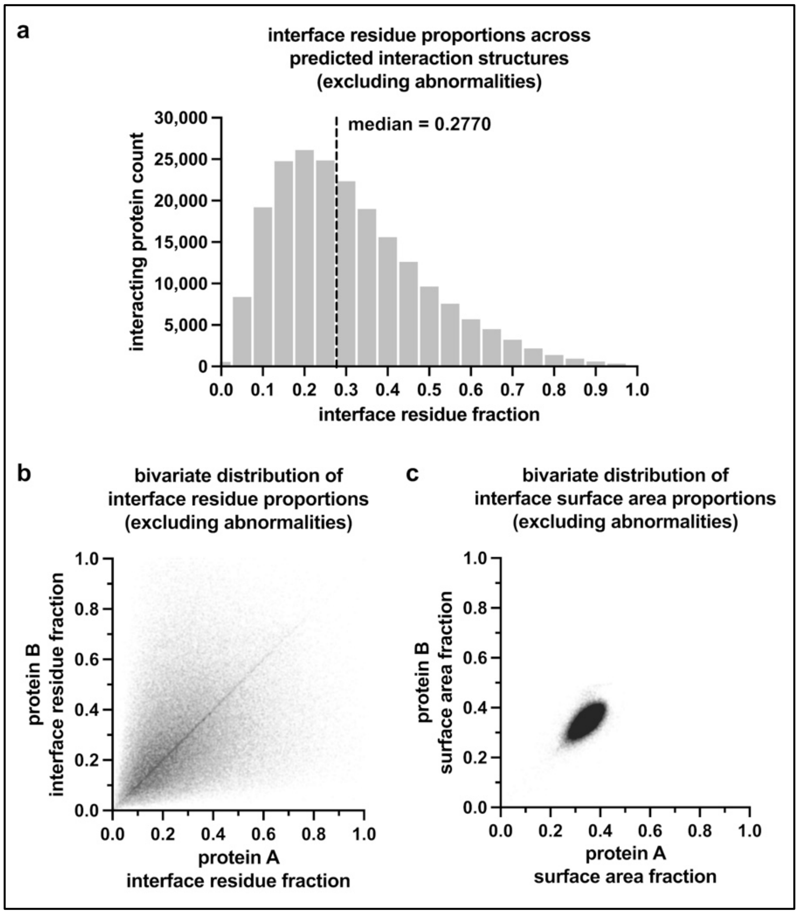


**Supplementary Fig. 2: Predicted structural protein interaction interfaces excluding structures with abnormalities. a.** Histogram of interface residue fractions for protein interaction structures excluding structures with no detectable interface and 10% structural overlap (as described in the main text). Interface residue fractions for proteins A and B were pooled into a single distribution. The median interface residue fraction (0.2770) is indicated by the dashed line. Histogram bin width = 0.05; $n$ = 209,494 interaction interface observations from 104,747 interaction structures. **b.** Bivariate distribution of interface residue fractions between the two interacting proteins in predicted protein interaction structures excluding structures with no detectable interface and 10% structural overlap. Each point represents one unique interaction. Statistical analysis by Spearman correlation ($r_S$ = 0.4016, 95% CI 0.3964 to 0.4068, $p < 0.0001$) and Pearson correlation ($r$ = 0.3393, 95% CI 0.3339 to 0.3446, $p < 0.0001$), $n$ = 104,747 interaction structures. **c.** Bivariate distribution of interface surface area fractions between the two interacting proteins in predicted protein interaction structures excluding structures with no detectable interface and 10% structural overlap. Each point represents one unique interaction. Statistical analysis by Spearman correlation ($r_S$ = 0.6469, 95% CI 0.6433 to 0.6505, $p < 0.0001$) and Pearson correlation ($r$ = 0.6626, 95% CI 0.6592 to 0.6660, $p < 0.0001$), $n$ = 104,747 interaction structures.

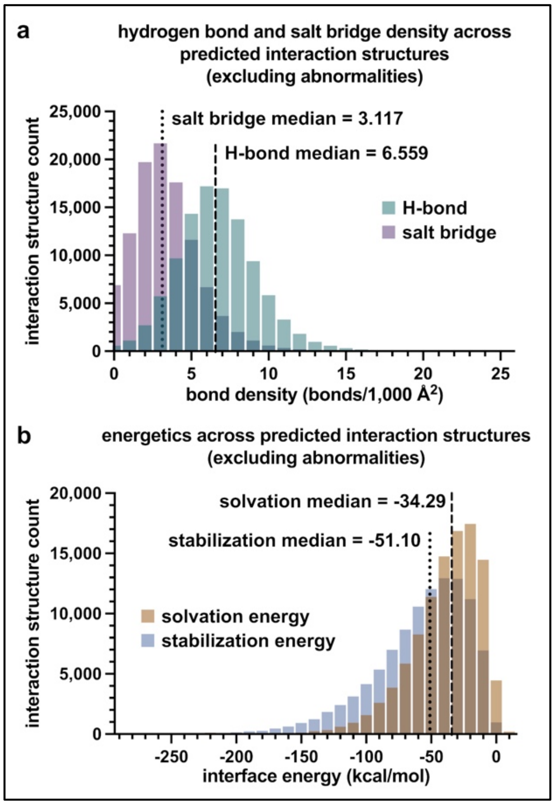


**Supplementary Fig. 3: Bonding and energetic properties excluding structures with abnormalities. a.** Histogram of intermolecular hydrogen bond and salt bridge densities for protein interaction structures excluding structures with no detectable interface and 10% structural overlap (as described in the main text). Hydrogen bond and salt bridge densities are displayed as separate distributions, distinguished by color. Bond densities are quantified as bonds per 1,000 Å² of mean BSA, as specified in **equation 4**. Medians are indicated by the dashed and dotted lines for hydrogen bond density (6.559) and salt bridge density (3.117), respectively. Histogram bin width = 1.0; *n* = 104,747 interaction structures. The *x*-axis was restricted to the range used in **Fig. 4a** for visual interpretability and comparability; all *n* = 104,747 data points, including those above this threshold, were retained for statistical analysis. **d.** Histogram of solvation and stabilization energies for protein interaction structures excluding structures with no detectable interface and 10% structural overlap, displayed as separate distributions distinguished by color. Medians are indicated by the dashed and dotted lines for solvation energy (-34.29) and stabilization energy (-51.10), respectively. Histogram bin width = 10; *n* = 104,747 interaction structures. The *x*-axis was restricted to the range used in **Fig. 4d** for visual interpretability and comparability; all *n* = 104,747 data points, including those above this threshold, were retained for statistical analysis.

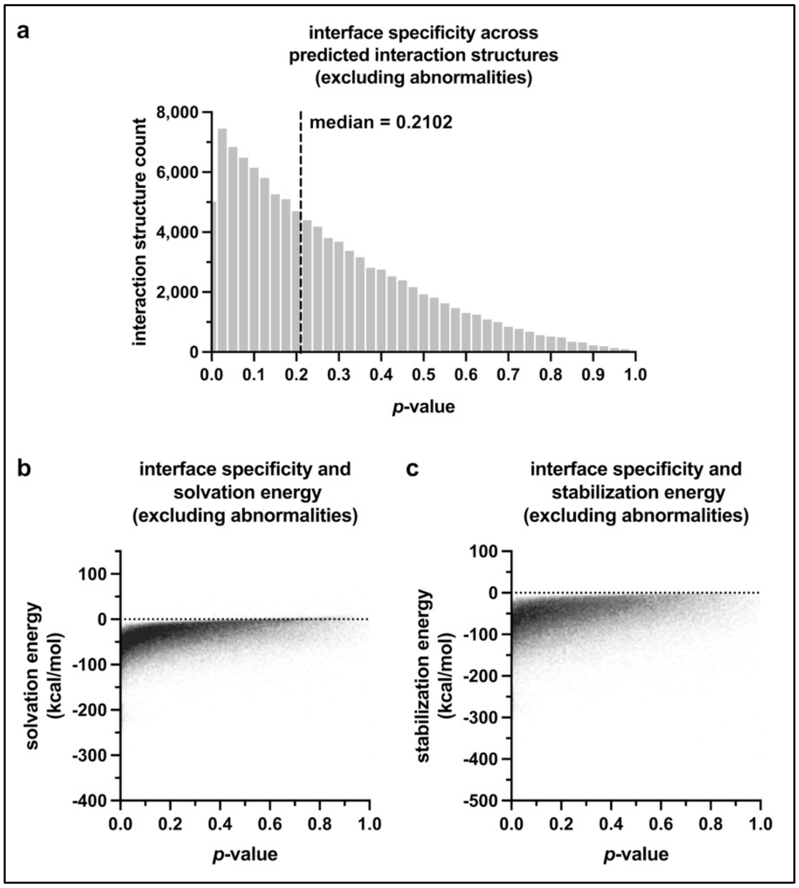


**Supplementary Fig. 4: Interface specificity and energetic trends excluding structures with abnormalities. a.** Histogram of the PISA-calculated *p*-values for interacting protein structures excluding structures with no detectable interface and 10% structural overlap (as described in the main text). The median p-value (0.2102) is indicated by the dashed line. Histogram bin width = 0.025; $n$ = 104,747 interaction structures. **b-c**. Relationships between *p*-value and **(f)** solvation energy or **(g)** stabilization energy for protein interaction structures excluding structures with no detectable interface and 10% structural overlap (as described in the main text). Each point represents one unique interaction. Statistical analysis by Spearman correlation (solvation: $r_S$ = 0.4858, 95% CI 0.4810 to 0.4905, $p < 0.0001$; stabilization: $r_S$ = 0.3777, 95% CI 0.3724 to 0.3831, $p < 0.0001$) and Pearson correlation (solvation: $r$ = 0.4102, CI 0.4051 to 0.4152, $p < 0.0001$; stabilization: $r$ = 0.3273, CI 0.3219 to 0.3327, $p < 0.0001$), $n$ = 104,747 interaction structures.
